# Full Brain and Lung Prophylaxis against SARS-CoV-2 by Intranasal Lentiviral Vaccination in a New hACE2 Transgenic Mouse Model or Golden Hamsters

**DOI:** 10.1101/2021.02.03.429211

**Authors:** Min-Wen Ku, Pierre Authié, Maryline Bourgine, François Anna, Amandine Noirat, Fanny Moncoq, Benjamin Vesin, Fabien Nevo, Jodie Lopez, Philippe Souque, Catherine Blanc, Sébastien Chardenoux, llta Lafosse, David Hardy, Kirill Nemirov, Françoise Guinet, Francina Langa Vives, Laleh Majlessi, Pierre Charneau

## Abstract

Non-integrative, non-cytopathic and non-inflammatory lentiviral vectors are particularly suitable for mucosal vaccination and recently emerge as a promising strategy to elicit sterilizing prophylaxis against SARS-CoV-2 in preclinical animal models. Here, we demonstrate that a single intranasal administration of a lentiviral vector encoding a prefusion form of SARS-CoV-2 spike glycoprotein induces full protection of respiratory tracts and totally avoids pulmonary inflammation in the susceptible hamster model. More importantly, we generated a new transgenic mouse strain, expressing the human Angiotensin Converting Enzyme 2, with unprecedent brain permissibility to SARS-CoV-2 replication and developing a lethal disease in <4 days post infection. Even though the neurotropism of SARS-CoV-2 is now well established, so far other vaccine strategies under development have not taken into the account the protection of central nervous system. Using our highly stringent transgenic model, we demonstrated that an intranasal booster immunization with the developed lentiviral vaccine candidate achieves full protection of both respiratory tracts and brain against SARS-CoV-2.

## Introduction

Severe Acute Respiratory Syndrome beta-coronavirus 2 (SARS-CoV-2) is the causative agent of pandemic coronavirus disease 2019 (COVID-19), hence the imminent necessity to develop effective and safe prophylactic vaccines. We have recently established the high performance of lentiviral vector (LV)-based vaccination against SARS-CoV-2 in pre-clinical animal models (Ku et al., 2021). Our reported COVID-19 LV-based vaccine candidate is currently in progress to proceed to clinical trial. Both integrative (ILV) and non-integrative (NILV) forms of LV allow transgene insertion up to 8kb in length and offer outstanding potential of gene transfer to the nuclei of host cells (Di Nunzio et al., 2012; Hu et al., 2011; Ku et al., 2020; Zennou et al., 2000). LVs display *in vivo* tropism for immune cells, notably dendritic cells (Arce et al., 2009, our unpublished data). They are non-replicative, non-cytopathic and scarcely inflammatory (unpublished data). These vectors induce long-lasting B- and T-cell immunity (Di Nunzio et al., 2012; Hu et al., 2011; Ku et al., 2020; Zennou et al., 2000). LV are pseudo-typed with the surface glycoprotein of Vesicular Stomatitis Virus, to which the human population has limited exposure, avoiding these vectors to be targeted by preexisting immunity in humans, which is in net contrast to adenoviral vectors (Rosenberg et al., 1998; Schirmbeck et al., 2008). The safety of LV has been established in human in a phase I/II Human Immunodeficiency Virus (HIV)-1 vaccine trial (2011-006260-52 EN).

“LV:: S_FL_” encodes the full-length sequence of the surface class I fusion spike (S) glycoprotein of SARS-CoV-2 (S_CoV-2_) that is able to elicit strong neutralizing antibodies (NAbs) and CD8^+^ T-cell responses against a large spectrum of S_CoV-2_ MHC-I-restricted epitopes (Ku et al., 2021). Notably, a systemic prime, followed by intranasal (i.n.) boost with LV::S_FL_ is associated with a robust prophylactic effect against lung SARS-CoV-2 replication, accompanied by strong reduction of infection-derived lung inflammation and tissue injury. These observations have been made in mice in which the expression of human Angiotensin-Converting Enzyme 2 (hACE2) was induced in the respiratory tracts by instillation of an adenoviral vector serotype 5 (Ad5::hACE2), or in the highly susceptible *Mesocricetus auratus*, i.e., outbred golden hamsters. These results indicated the requirement for i.n. boost and induction of mucosal adaptive immunity to reach almost sterilizing lung protection against SARS-CoV-2 (Ku et al., 2021).

Although lung is the organ of predilection for SARS-CoV-2, its neurotropism, similar to that of SARS-CoV and Middle East Respiratory Syndrome (MERS)-CoV, (Glass et al., 2004; Li et al., 2016; Netland et al., 2008) has been reported (Aghagoli et al., 2020; Fotuhi et al., 2020; Hu et al., 2020; Liu et al., 2020; Politi et al., 2020; Roman et al., 2020; von Weyhern et al., 2020; Whittaker et al., 2020). Moreover, expression of ACE2 in neuronal and glial cells has been described (Chen et al., 2020; Xu and Lazartigues, 2020). Accordingly, COVID-19 human patients can present symptoms like headache, myalgia, anosmia, dysgeusia, impaired consciousness and acute cerebrovascular disease (Bourgonje et al., 2020; Hu et al., 2020; Mao et al., 2020). Analysis of autopsies of COVID-19 deceased patients demonstrated presence of SARS-CoV-2 in nasopharynx and brain, and virus entry into central nervous system (CNS) via neural-mucosal interface of olfactory mucosa (Meinhardt et al., 2020). These observations align with the previous report of viruses capable of gaining access to the brain through neural dissemination or hematogenous route (Desforges et al., 2014). Therefore, it is critical to focus hereinafter on the protective properties of COVID-19 vaccine candidates, not only in the respiratory tracts, but also in the brain.

Contribution of S_CoV-2_ is instrumental in host cell invasion through cellular attachment and membrane fusion subsequent to engagement with hACE2 (Walls et al., 2020). Following this interaction, the extracellular domain of the metastable prefusion S_CoV-2_ is cleaved at the 682^RRAR^685 site of subtilisin-like protease furin (Guo et al., 2020; Walls et al., 2020). This gives rise to: (i) S1 subunit, harboring the ACE2 Receptor Binding Domain (RBD) that harbors main B-cell epitopes targeted by NAbs (Walls et al., 2020), and (ii) S2 subunit, containing the membrane-fusion machinery. This cleavage leads to substantial conformational rearrangements resulting in a highly stable post-fusion form of S_CoV-2_ that initiates the fusion reaction with the host cell membrane (Sternberg and Naujokat, 2020). Similar to envelop glycoproteins of several other viruses that harbor furin cleavage site, including respiratory syncytial virus, HIV, Ebola virus, human metapneumovirus, and Lassa virus (Bos et al., 2020). It is possible to engineer S_CoV-2_ to limit its conformational dynamics and avoid the stabilization under its prefusion conformation to maintain better exposure of the S1 B-cell epitopes and improve immunogen availability (McCallum et al., 2020).

Here, we first demonstrated that a single i.n. administration of an LV encoding a prefusion form of S_CoV-2_ was as protective as a systemic prime followed by an i.n. boost by conferring sterilizing pulmonary protection against SARS-CoV-2 in the pre-clinical hamster model. A single i.n. injection of this LV was as protective as a systemic prime followed by an i.n. boost in these animals. More importantly, we generated a new hACE2 transgenic mouse strain with unprecedent permissibility of the brain to SARS-CoV-2 replication. Using this unique preclinical animal model, we demonstrated the importance of i.n. booster immunization with the developed LV-based vaccine candidate to reach full protection of not only lungs but also CNS against SARS-CoV-2. Our results indicated that i.n. vaccination with non-cytopathic and non-inflammatory LV is a performant and safe strategy to elicit sterilizing immunity in the main anatomical sites affected by COVID-19.

## Results

### Adaptive Immunity Elicited by LV::S_ΔF2P_ in Comparison to LV::S_FL_

We generated LV encoding a prefusion form of S_CoV-2_ under transcriptional control of the cytomegalovirus promoter. This prefusion S_CoV-2_ variant (S_ΔF2P_) has a deletion of 675^QTQTNSPRRAR^685 sequence. This deletion encompasses the polybasic RRAR furin cleavage site situated at the junction of S1/S2 subunits, and harbors K^986^P and V^987^P consecutive proline substitutions in S2, within the hinge loop between heptad repeat 1 and the central helix (Figure S1A) (Walls et al., 2020). Both ILV::S_ΔF2P_- and ILV::S_FL_-primed (i.m.) and -boosted (i.n.) C57BL/6 mice possessed high serum titers of anti-S_CoV-2_ IgG (Figure S1B) and high titers of anti-S_CoV-2_ IgG and IgA in their lung extracts (Figure S1C), indicating that the modifications in the pre-fusion S_ΔF2P_ form does not impact positively or negatively its capacity to induce Abs against native S_CoV-2_.

### Sterilizing protection in hamster model by a single i.n. NILV::S_ΔF2P_ administration

We recently showed that despite induction of high serum neutralizing activity, systemic immunization with ILV::S_FL_ or NILV::S_FL_ conferred significant but partial protection against SARS-CoV-2. In contrast, vaccination by systemic prime followed by i.n. boost with ILV::S_FL_ or NILV::S_FL_ induced almost sterilizing lung protection (Ku et al., 2021). Here, we first assessed the prophylactic effect of vaccination with only a single i.n. administration of NILV::S_ΔF2P_ in the hamster model.

Hamsters (*n* = 6/group) were: (i) primed i.m. at wk 0 with 1 × 10^8^ TU of NILV::S_ΔF2P_ and boosted i.n. at wk 5 with the same amount of the vector, as a positive protection control, (ii) immunized i.n. with a single injection of 1 × 10^8^ TU of NILV::S_ΔF2P_ at wk 0, or (iii) at wk 5 (Figure 1A). Sham-vaccinated controls received equivalent amounts of an NILV::GFP via i.n. at wks 0 and 5. Comparable and high titers of anti-S_CoV-2_ IgG Abs were detected in the sera in the first two groups at wk 5 (Figure 1B). At wk 7, the serum Ab titer was maintained at a high level in the NILV:: S_ΔF2P_ i.m.-i.n. group while it was slightly decreased in some individuals of the “NILV::S_ΔF2P_ i.n. wk 0” group. At this time point, in the “NILV::S_ΔF2P_ i.n. wk 5” group, lower serum Ab titers were detected (Figure 1B). Although the virus neutralization activity was significantly lower in the sera of “NILV::S_ΔF2P_ i.n. wk 5” hamsters compared to the NILV::S_ΔF2P_ i.m.-i.n. group, the former had an equivalent neutralizing capacity in their lung homogenates (Figure 1C).

**Figure 1.**
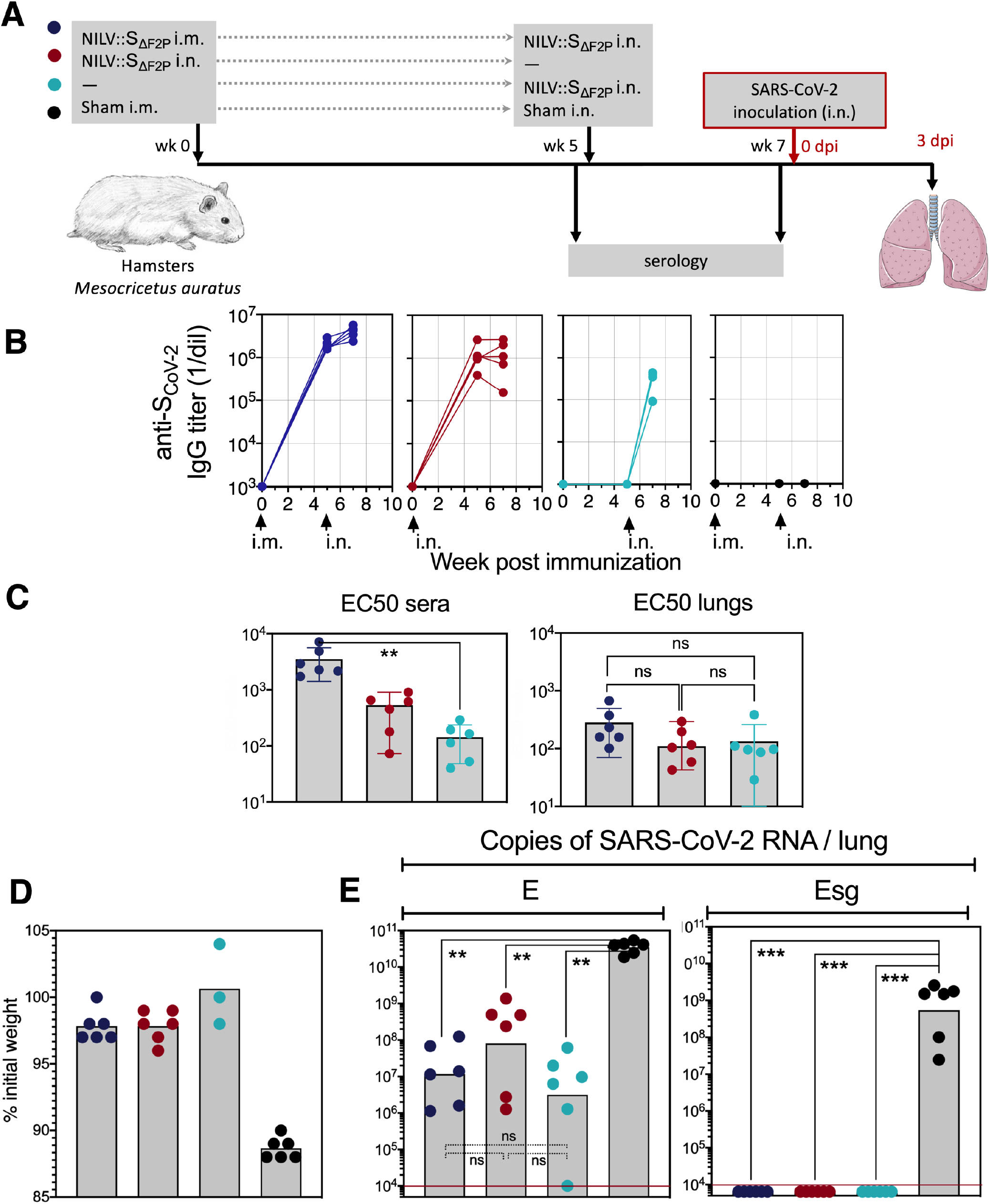
Single i.n. injection of LV::S_ΔF2P_ fully protects golden hamsters against SARS-CoV-2. **(A)** Timeline of the LV::S_ΔF2P_ prime-boost vaccination regimen and SARS-CoV-2 challenge in hamsters. **(B)** Serum anti-S_CoV-2_ IgG responses expressed as mean endpoint dilution titers, determined by ELISA. **(C)** Neutralization capacity of anti-S_CoV-2_ Abs, expressed as EC50 neutralizing titers, determined in the sera and lung homogenates of LV:: S_ΔF2P_-immunized hamsters. Bars represent mean ± SD. **(D)** Percentages of weight loss in LV::S_ΔF2P_- or sham-vaccinated hamsters at 4 dpi. **(E)** Lung viral loads quantitated by total E or Esg qRT-PCR at 4 dpi. Statistical significance of the differences was evaluated by two tailed unpaired t test; ** = *p*<0.01, *** = *p* <0.001. Red lines indicate the limit of detection of each assay.

At wk 7, all animals were challenged i.n. with 0.3 × 10^5^ TCID50 of a SARS-CoV-2. At 4 days post inoculation (dpi), only 2-3% weight loss was detected in the NILV::S_ΔF2P_-vaccinated groups, compared to 12% in sham-vaccinated hamsters (Figure 1D). At this time point, as determined by qRT-PCR detecting SARS-CoV-2 Envelop (E_CoV-2_) RNA, ~ 2-to-3 log10 decreases were observed in NILV::S_ΔF2P_-vaccinated individuals of either i.m.-i.n. or single i.n. groups, compared to sham-vaccinated group (Figure 1E).

Assessment of lung viral load by a sub-genomic E_CoV-2_ RNA (Esg) qRT-PCR, which was reported to be an indicator of active viral replication (Chandrashekar et al., 2020; Tostanoski et al., 2020; Wolfel et al., 2020) (Tostanoski et al., 2020), showed total absence of replicating virus in the three vaccinated groups versus a mean ± SD of (1.24 ± 0.99) × 10^9^ copies of Esg RNA of SARS-CoV-2/lungs in the sham-vaccinated group (Figure 1E). Many publications use PFU counting to determine viral loads. We noticed that large amounts of NAbs in the lungs of vaccinated individuals, even though not necessarily spatially in contact with circulating viral particles in alive animal, can come to contact with and neutralize viral particles in the lung homogenates in vitro. In this case the PFU assay underestimates the amounts of cultivable viral particles (Figure S2A).

At 4 dpi, as evaluated by qRT-PCR in total lung homogenates, a substantial decreased in inflammation was detected in NILV::S_ΔF2P_-vaccinated hamsters compared to their sham-vaccinated counterparts, regardless of the immunization regimen, i.e., i.m.-i.n. prime-boost or single i.n. injection given at wk 0 or 5 (Figure 2A). On lung histopathological examination, sham-vaccinated controls demonstrated lung infiltration and interstitial syndrome (Figure 2B-D), severe alveolo-interstitial inflammation leading to dense pre-consolidation areas (Figure 2E), accompanied by bronchiolar lesions, with images of bronchiolar epithelium desquamation into the lumen (Figure 2F-H). In vaccinated groups the lesions were minimal, although some degree of alveolar infiltration could be seen (Figure 2I).

**Figure 2.**
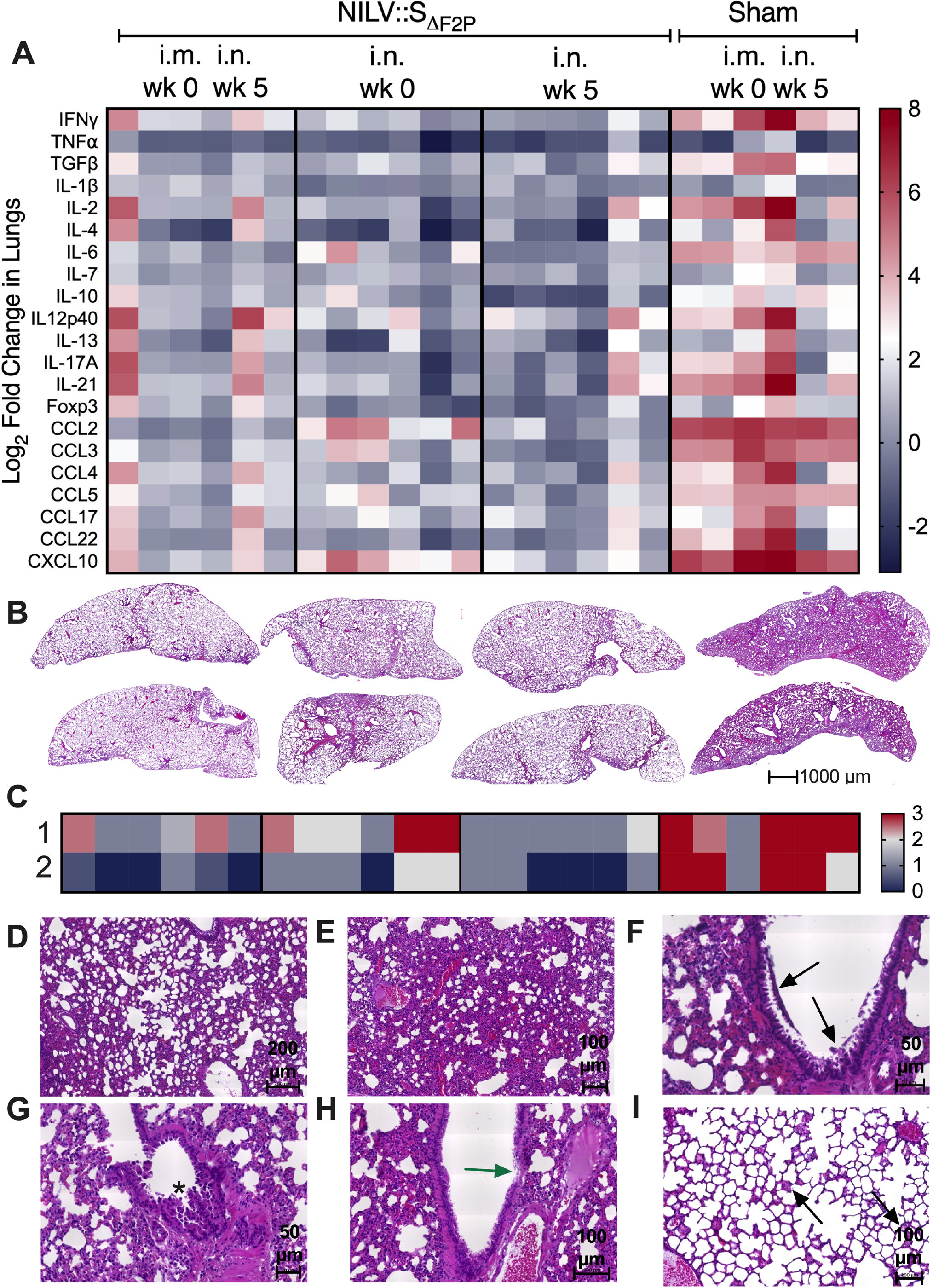
Largely reduced infection-driven lung inflammation in LV::S_ΔF2P_-vaccinated hamsters. **(A)** Heatmap recapitulating relative log2 fold changes in the expression of inflammation-related mediators in S_ΔF2P_- or sham-vaccinated individuals, as analyzed at 4 dpi by use of RNA extracted from total lung homogenates and normalized versus samples from untreated controls. Six individual hamsters per group are shown in the heatmap. **(B)** Lung histological H&E analysis, as studied at 4 dpi. **(C)** Heatmap recapitulating the histological scores, for: 1) inflammation score and 2) interstitial syndrome. **(D)** Representative alveolo-interstitial syndrome and **(E)** severe inflammation in a sham-vaccinated and infected hamster. Here the structure of the organ is largely obliterated, while remnants of alveolar spaces and bronchiolar lumens can be seen. (**F-H**) bronchiolar lesions in sham-vaccinated animals. Shown are epithelial cells and cell debris in the bronchiolar lumen (black arrows) **(F),** papillary projections of the bronchiolar epithelium into the lumen (star) (**G**) and degenerative lesions with effacement of the epithelium (green arrow) (**H**). **(I)** Mild alveolar infiltration in a vaccinated hamster. Some of the alveoli (arrow) are partially or completely filled with cells and an eosinophilic exudate.

These data collectively indicated that a single i.n. administration of NILV::S_ΔF2P_ was as protective as a systemic prime and i.n. boost regimen, conferred sterilizing pulmonary immunity against SARS-CoV-2 and readily prevented lung inflammation and pathogenic tissue injury in the susceptible hamster model. These data also showed the long-term feature of the conferred protection because 7 weeks after a single injection of the vaccine, the protection potential remained complete.

### Generation of new hACE2 transgenic mice with substantial brain permissibility to SARS-CoV-2 replication

No hACE2 transgenic mice were available in Europe until September 2020. To set up a mouse model permissive to SARS-CoV-2 replication allowing assessment of our vaccine candidates, based on the previously produced B6.K18-ACE2^2Prlmn/JAX^ mice (McCray et al., 2007), we generated C57BL/6 transgenic mice with an LV (Nakagawa and Hoogenraad, 2011) carrying the *hACE2* gene under the human cytokeratin 18 promoter, namely “B6.K18-hACE2^IP-THV^”. The permissibility of these mice to SARS-CoV-2 replication was evaluated after one generation backcross to WT C57BL/6 (N1). N1 mice with varying number of *hACE2* transgene copies per genome (Figure 3A) were sampled and inoculated i.n. with SARS-CoV-2 (Figure 3B). At 3 dpi, the mean ± SD of lung viral loads was as high as (3.3 ± 1.6) × 10^10^ copies of SARS-CoV-2 RNA/lung in permissive mice (Figure 3B). SARS-CoV-2 RNA copies per lung <1 × 10^7^ correspond to the genetic material derived from the input in the absence of viral replication (Ku et al., 2021). We also noted that the lung viral loads (Figure 3B) were not proportional to the *hACE2* transgene copy number per genome (Figure 3A). Remarkably, substantial viral loads, i.e., (5.7 ± 7.1) × 10^10^ copies of SARS-CoV-2 RNA, were also detected in the brain of the permissive mice (Figure 3B). Virus replication/dissemination was also observed, although to a lower extent, in the heart and kidneys.

**Figure 3.**
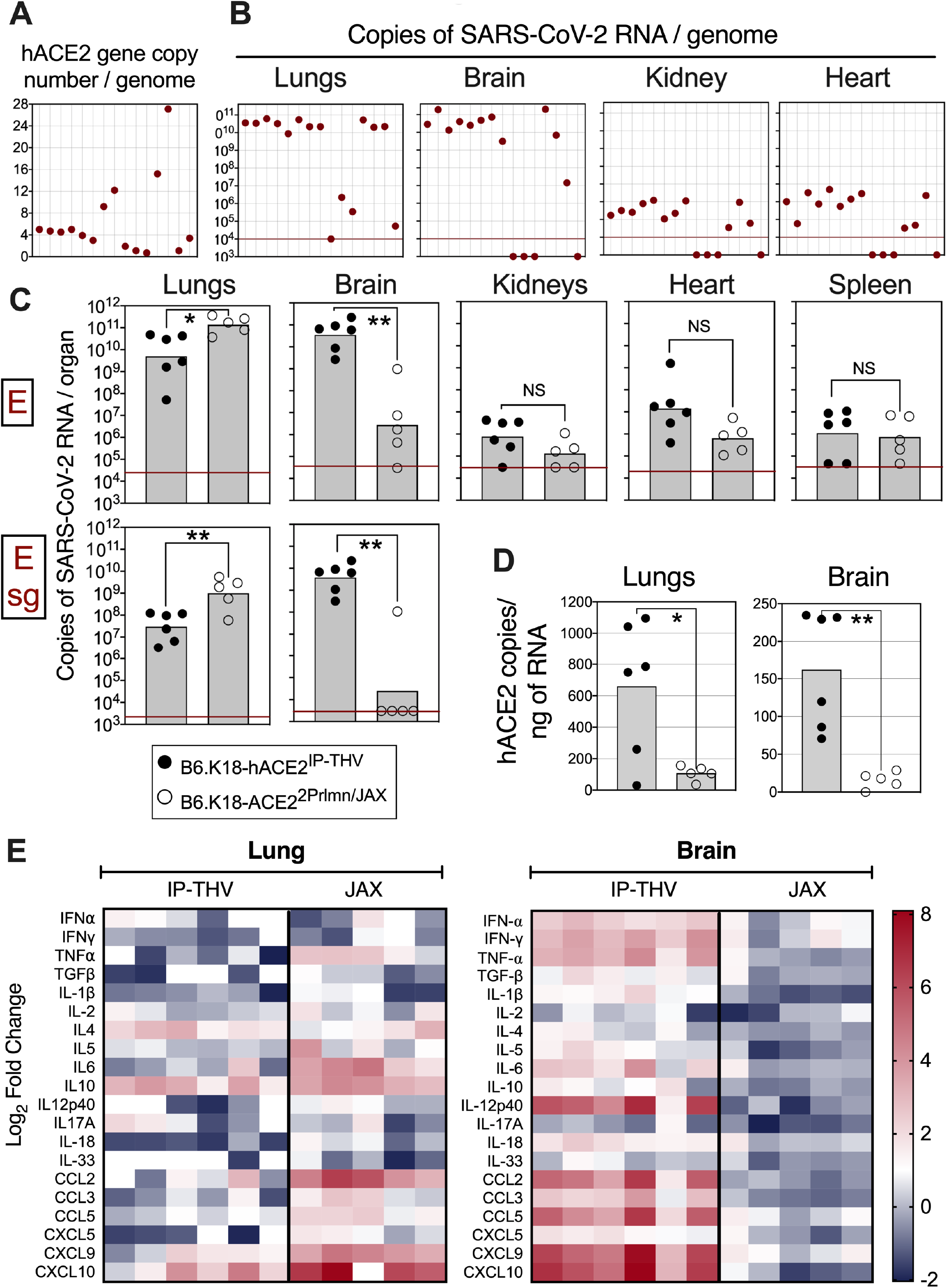
Large permissibility of the lungs and brain of K18-hACE2^IP-THV^ transgenic mice to SARS-CoV-2 replication. **(A)** Representative genotyping results from 15 N1 B6.K18-hACE2^IP-THV^ mice as performed by qPCR to determine their *hACE2* gene copy number per genome. Dots represent individual mice. **(B)** Phenotyping of the same mice, inoculated i.n. with 0.3 × 10^5^ TCID50 at the age of 5-7 wks and viral loads determination in their various organs at 3 dpi by conventional E-specific qRT-PCR. **(C)** Comparative permissibility of diverse organs from K18-hACE2^IP-THV^ and B6.K18-ACE2^2Prlmn/JAX^ transgenic mice to SARS-CoV-2 replication, as determined at 3 dpi by conventional E-specific or sub-genomic Esg-specific qRT-PCR. Red lines indicate the qRT-PCR limits of detection. Statistical significance of the difference was evaluated by Mann-Whitney test (*= *p* < 0.01, **= *p* <0.00). **(D)** Comparative quantitation of *hACE-2* mRNA in the lungs and brain of B6.K18-hACE2^IP-THV^ and B6.K18-ACE2^2Prlmn/JAX^ transgenic mice. **(E)** Heatmap recapitulating log2 fold change in cytokine and chemokine mRNA expression in the lungs or brain of B6.K18-hACE2^IP-THV^ and B6.K18-ACE2^2Prlmn/JAX^ transgenic mice at 3 dpi. Data were normalized versus untreated controls. Statistical significance of the difference was evaluated by Mann-Whitney test (*=*p* < 0.05, **=*p* <0.01).

We further compared the replication of SARS-CoV-2 in lungs and brain and the viral dissemination to various organs in B6.K18-hACE2^IP-THV^ and B6.K18-ACE2^2Prlmn/JAX^ mice (McCray et al., 2007) (Figure 3C). The lung viral loads were slightly lower in B6.K18-hACE2^IP-THV^ compared to B6.K18-ACE2^2Prlmn/JAX^ mice. However, viral loads in the brain of B6.K18-hACE2^IP-THV^ mice were substantially higher compared to their B6.K18-ACE2^2Prlmn/JAX^ counterparts (Figure 3C). Measurement of brain viral loads by Esg qRT-PCR detected (7.55 ± 7.74) × 10^9^ copies of SARS-CoV-2 RNA in B6.K18-hACE2^IP-THV^ mice and no copies of this replication-related RNA in 4 out of 5 B6.K18-ACE2^2Prlmn/JAX^ mice. This substantial difference of SARS-CoV-2 replication in the brain of both transgenic strains was corroborated with significantly higher *hACE2* mRNA expression in the brain of B6.K18-hACE2^IP-THV^ mice (Figure 3D). However, *hACE2* mRNA expression in the lungs of B6.K18-hACE2^IP-THV^ mice was also higher than in B6.K18-ACE2^2Prlmn/JAX^ mice, which does not account for the lower viral replication in the lungs of the former. A trend towards higher viral loads was also observed in the kidneys and heart of B6.K18-hACE2^IP-ThV^ compared to B6.K18-ACE2^2Prlmn/JAX^ mice (Figure 3C).

In concordance with the lung viral loads, as evaluated by qRT-PCR applied to total lung homogenates, B6.K18-hACE2^IP-THV^ mice displayed less pulmonary inflammation than B6.K18-ACE2^2Prlmn/JAX^ mice (Figure 3E). Remarkably, this assay applied to total brain homogenates detected substantial degrees of inflammation in B6.K18-hACE2^IP-THV^ — but not in B6.K18-ACE2^2Prlmn/JAX^ — mice (Figure 3E). In addition, B6.K18-hACE2^IP-THV^ mice reached the humane endpoint between 3 and 4 dpi and therefore displayed a lethal SARS-CoV-2-mediated disease more rapidly than their B6.K18-ACE2^2Prlmn/JAX^ counterparts (Winkler et al., 2020).

Therefore, the large permissibility to SARS-CoV-2 replication at both lung and CNS, marked brain inflammation and rapid development of a lethal disease are major distinctive features offered by this new B6.K18-hACE2^IP-THV^ transgenic model.

### Full protection of lungs and brain in LV::S_ΔF2P_-immunized B6.K18-hACE2^IP-THV^ mice

We then evaluated the vaccine efficacy of LV::S_ΔF2P_ in B6.K18-hACE2^IP-THV^ mice. In a first experiment with these mice, we used an integrative version of the vector. Individuals (*n* = 6/group) were primed i.m. with 1 × 10^7^ TU/mouse of ILV::S_ΔF2P_ or an empty LV (sham) at wk 0 and then boosted i.n. at wk 3 with the same dose of the same vectors (Figure 4A). Mice were then challenged with SARS-CoV-2 at wk 5. A high serum neutralizing activity, i.e., EC50 mean ± SD of 5466 ± 6792, was detected in ILV::S_ΔF2P_-vaccinated mice Figure 4B). This vaccination conferred substantial degrees of protection against SARS-CoV-2 replication, not only in the lungs, but also in the brain (Figure 4C). Notably, quantitation of brain viral loads by Esg qRT-PCR detected no copies of this replication-related SARS-CoV-2 RNA in ILV::S_ΔF2P_-vaccinated mice *versus* (7.55 ± 7.84) × 10^9^ copies in the brain of the sham-vaccinated controls.

**Figure 4.**
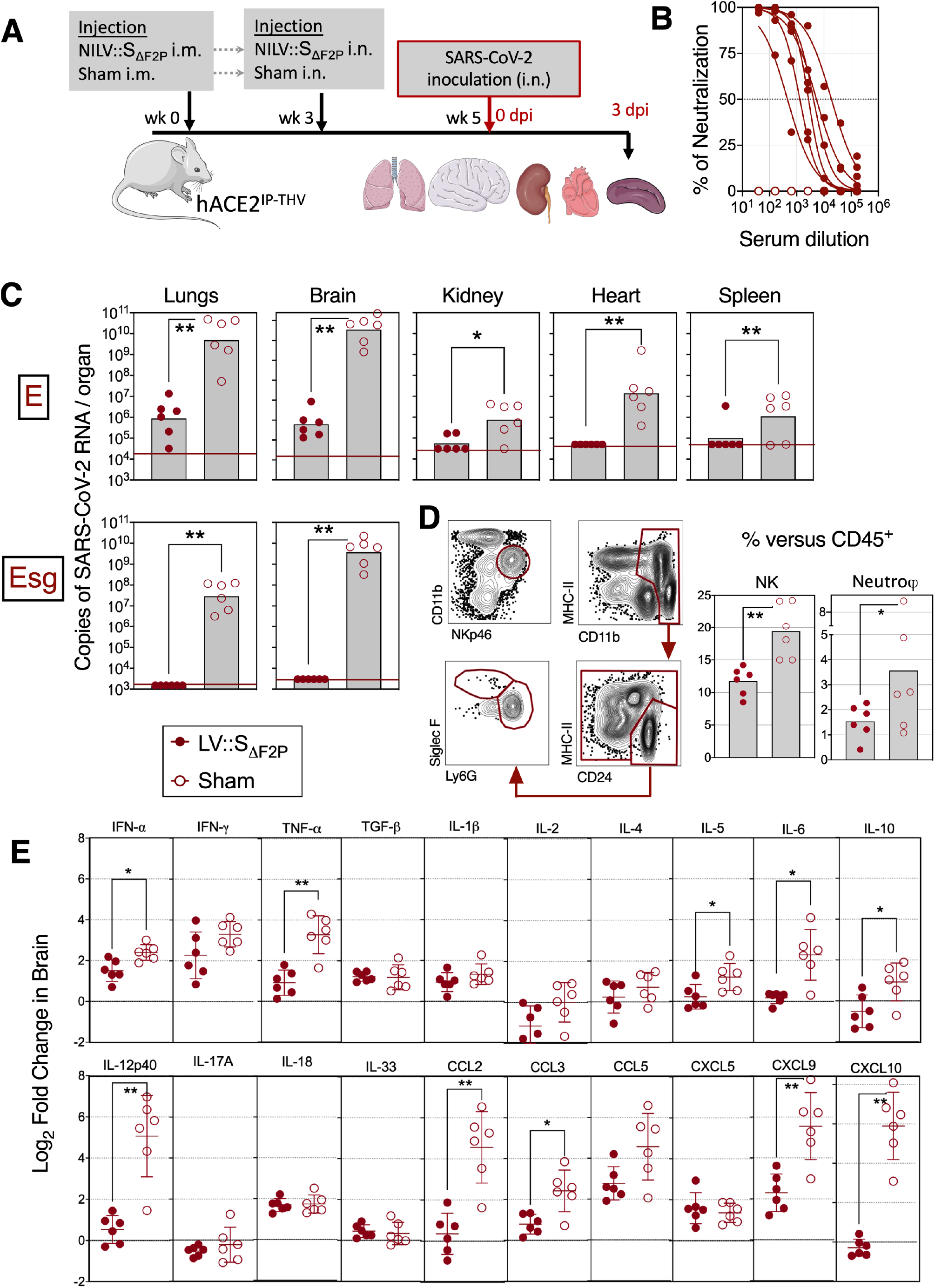
Vaccination with LV::S_ΔF2P_ protects both lungs and central nervous system from SARS-CoV-2 infection in K18-hACE2^IP-THV^ transgenic mice. **(A)** Timeline of prime-boost LV::S_ΔF2P_ vaccination and SARS-CoV-2 challenge in K18-hACE2^IP-THV^ mice. **(B)** Serum neutralization capacity of anti-S_CoV-2_ Abs in LV::S_ΔF2P_-vaccinated mice. **(C)** Viral loads as determined in diverse organs at 3dpi by use of conventional E-specific or sub-genomic Esg-specific qRT-PCR. Red lines indicate the qRT-PCR detection limits. Statistical significance of the difference was evaluated by Mann-Whitney test (*= *p* < 0.01, **= *p* <0.001). **(D)** Cytometric gating strategy determined to identify and quantify lung NK cells and neutrophils in the lungs of LV::S_ΔF2P_- or sham-vaccinated and SARS-CoV-2-challenged K18-hACE2^IP-THV^ transgenic mice at 3 dpi. Percentages of NK and neutrophil subset were calculated among total lung CD45^+^ cells. **(E)** Relative log2 fold change in cytokine and chemokine mRNA expression in the brain of LV:: S_ΔF2P_- or sham-immunized and SARS-CoV-2-challenged K18-hACE2^IP-THV^ transgenic mice at 3 dpi. Data were normalized versus untreated controls. Statistical significance was evaluated by two tailed unpaired t test; * = p<0.05, ** = p<0.01).

At 3 dpi, cytometric investigation of the lung innate immune cell subsets (Figure 4D, S3A) detected significant decrease in the proportions of NK (CD11b^int^ NKp46^+^) cells and neutrophils (CD11b^+^ CD24^+^ SiglecF^-^ Ly6G^+^) among the lung CD45^+^ cells in the ILV::S_ΔF2P_-vaccinated and protected B6.K18-hACE2^IP-THV^ mice, compared to the sham-vaccinated and unprotected controls (Figure 4D). Frequencies of the other lung innate immune cells were not significantly distinct in the protected and unprotected groups (Figure S3B). At 3 dpi, as evaluated by qRT-PCR applied to brain homogenates, ILV::S_ΔF2P_-vaccinated B6.K18-hACE2^IP-THV^ mice had significant decreases in the expression levels of IFN-a, TNF-a, IL-5, IL-6, IL-10, IL-12p40, CCL2, CCL3, CXCL9 and CXCL10, compared to the sham group (Figure 4E). Furthermore, no noticeable changes in the lung inflammation were recorded between the two groups (Figure S3C).

Therefore, an i.m.-i.n. prime-boost with ILV::S_ΔF2P_ prevents SARS-CoV-2 replication in both lung and CNS anatomical areas and inhibits virus-mediated lung infiltration and pathology and neuro-inflammation.

### Requirement of i.n. boost for full protection of brain in B6.K18-hACE2^IP-THV^ mice

To go further in characterization of the protective properties of LV, in the following experiments in B6.K18-hACE2^IP-THV^ mice, similar to the hamster model, we used the safe and non-integrative version of LV. The observed protection of brain against SARS-CoV-2 may reflect the benefits of i.n. route of LV administration against this respiratory and neurotropic virus. To address this question, B6.K18-hACE2^IP-THV^ mice were vaccinated with NILV::S_ΔF2P_: (i) i.m. wk 0 and i.n. wk5, as a positive control, (ii) i.n. wk 0, or (iii) i.m. wk 5. Sham-vaccinated controls received i.n. an empty NILV at wks 0 and 5 (Figure 5A). Mice were then challenged with SARS-CoV-2 at wk 7 and viral loads were determined in the brain by E- or Esg-specific qRT-PCR at 3dpi (Figure 5B). In this highly stringent pre-clinical model, even performant, a single i.n. or i.m. injection of NILV::S_ΔF2P_ did not induce full protection in all animals of each group. Only i.m. prime followed by i.n. boost conferred full protection in all animals, showing the requirement of an i.n. boost to reach full protection of brain. The brain protection levels were proportional to plasma anti-S_CoV-2_ IgG titers (Figure 5C). As analyzed by cytometry, composition of innate and adaptive immune cells in the cervical lymph nodes were unchanged in NILV::S_ΔF2P_ i.m.-i.n. protected group, sham i.m.-i.n. unprotected group and untreated controls (data not shown). However, we detected increased proportion of CD8^+^ T cells in the olfactory bulb of NILV::S_ΔF2P_ i.m.-i.n. protected group compared to the sham unprotected group (Figure 5D). CD4^+^ T cells in the olfactory bulb had no distinctive activated or migratory phenotype, as assessed by their surface expression of CD69 or CCR7, respectively. In line with the absence of CCR7 expression on these T cells, and unlike the case of Murine Hepatitis Virus (MHV) infection (Cupovic et al., 2016), we did not detect up-regulation of CCL19 and CCL21 chemo-attractants (CCR7 ligands) in the brain and regardless of the mouse protection status (Figure 5E). Compared to NILV::S_ΔF2P_ i.m.-i.n. protected group, increased amount of neutrophils (CD11b^+^ Ly6C^+^ Ly6G^+^) in the olfactory bulb (Figure 5F) and inflammatory monocytes (CD11b^+^ Ly6C^+^ Ly6G^-^) in the brain (Figure 5G) of unprotected mice, compared to NILV::S_ΔF2P_ i.m.-i.n. protected group. The presence of these cells is a biomarker of inflammation and therefore, correlated with active viral replication.

**Figure 5.**
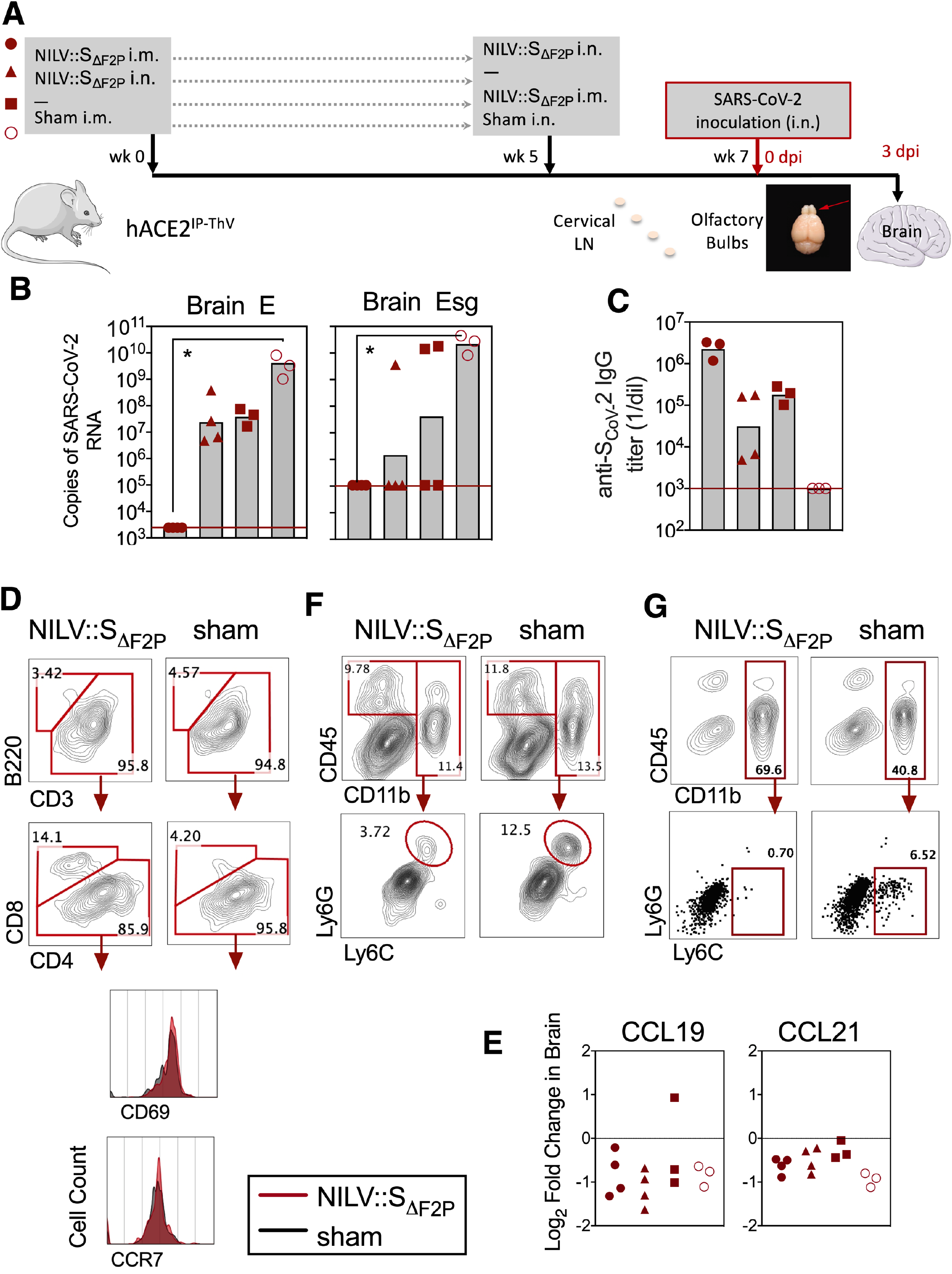
Booster vaccination with LV::S_ΔF2P_ through i.n. route elicits full protection of CNS from SARS-CoV-2 infection. **(A)** Timeline of various LV::S_ΔF2P_ vaccination regimens and SARS-CoV-2 challenge in B6.K18-hACE2^IP-THV^ mice. **(B)** Viral loads in the brain at 3dpi determined by conventional E-specific or sub-genomic Esg-specific qRT-PCR. Red lines indicate the qRT-PCR detection limits. Statistical significance of the difference was evaluated by Mann-Whitney test (*=*p* < 0.01). **(C)** Plasma anti-S_CoV-2_ Ab titers. **(D, F, G)** Cytometric analysis at 3 dpi performed on cells extracted from pooled olfactory bulbs or brain of LV::S_ΔF2P_ i.m.-i.n. vaccinated and protected mice versus sham-vaccinated and unprotected mice. **(D)** Adaptive and **(F)** innate immune cells in the olfactory bulbs. **(E)** Expression of CCL19 or CCL21 chemoattractant in the brain, as evaluated by qRT-PCR. **(G)** Innate immune cells in the brain.

Collectively, our data generated in the highly stringent B6.K18-hACE2^IP-THV^ mouse model support the advantage of NILV::S_ΔF2P_ i.n. boost in the immune protection of CNS from SARS-CoV-2 replication and the resulting infiltration and neuro-inflammation. Therefore, local induction and/or activation of mucosal immune response in the nasal cavity and olfactory bulbs, i.e. one of the potential entry points for the virus, emerges as a performant strategy.

## Discussion

LV-based platform emerges recently as a powerful vaccination approach against COVID-19, notably when used in systemic prime followed by mucosal i.n. boost, able to induce sterilizing immunity against lung SARS-CoV-2 infection in preclinical animal models (Ku et al., 2021). In the present study, we first demonstrated that a single i.n. administration of an LV encoding the S_ΔF2P_ prefusion form of S_CoV-2_ confers, as efficiently as an i.m. - i.n. prime-boost regimen, full protection of lungs in the highly susceptible hamster model, as evaluated by virological, immunological and histopathological parameters. The hamster ACE2 ortholog interacts efficaciously with S_CoV-2_, which readily allows host cell invasion by SARS-CoV-2 and its high replication rate. With rapid weight loss and development of severe lung pathology subsequent to SARS-CoV-2 inoculation, this species provides a sensitive model to evaluate the efficacy of drug or vaccine candidates, for instance compared to Rhesus macaques which develop only a mild COVID-19 pathology (Munoz-Fontela et al., 2020; Sia et al., 2020). The fact that a single i.n. LV administration, either seven or two weeks before SARS-CoV-2 challenge, elicits sterilizing lung protection in this susceptible model is valuable in setting the upcoming clinical trials with this LV-based vaccine and could provide remarkable socio-economic advantages for mass vaccination. The long-termed protection conferred, i.e., at least 7 weeks after a single injection, is another major advantage of this vaccine strategy to be translated in humans.

To further investigate the efficacy of our vaccine candidates, we generated a new transgenic mouse model, using the LV-based transgenesis approach (Nakagawa and Hoogenraad, 2011). The ILV used in this strategy encodes for hACE2 under the control of the cytokeratin K18 promoter, i.e., the same promoter as previously used by Perlman’s team to generate B6.K18-ACE2^2Prlmn/JAX^ mice (McCray et al., 2007), with a few adaptations to the lentiviral FLAP transfer plasmid. However, the new B6.K18-hACE2^IP-THV^ mice have certain distinctive features, as they express much higher levels of hACE2 mRNA in the brain and display markedly increased brain permissibility to SARS-CoV-2 replication, in parallel with a substantial brain inflammation and development of a lethal disease in <4 days post infection. These distinct characteristics can arise from differential hACE2 expression profile due to: (i) alternative insertion sites of ILV into the chromosome compared to naked DNA, and/or (ii) different effect of the Woodchuck Posttranscriptional Regulatory Element (WPRE) *versus* the alfalfa virus translational enhancer (McCray et al., 2007), in B6.K18-hACE2^IP-THV^ and B6.K18-ACE2^2Prlmn/JAX^ animals, respectively. Other reported *hACE2* humanized mice expressing the transgene under: (i) murine ACE2 promoter, without reported hACE2 mRNA expression in the brain (Yang et al., 2007), (ii) “hepatocyte nuclear factor-3/forkhead homologue 4” (HFH4) promoter, i.e., “HFH4-hACE2” C3B6 mice, in which lung is the principal site of infection and pathology (Jiang et al., 2020; Menachery et al., 2016), and (iii) “CAG” mixed promoter, i.e. “AC70” C3H × C57BL/6 mice, in which hACE2 mRNA is expressed in various organs including lungs and brain (Tseng et al., 2007). Comparison of AC70 and B6.K18-hACE2^IP-THV^ mice could yield information to assess the similarities and distinctions of these two models. However, here we report much higher brain permissibility of B6.K18-hACE2^IP-THV^ mice to SARS-CoV-2 replication, compared to B6.K18-ACE2^2Prlmn/JAX^ mice. The B6.K18-hACE2^IP-THV^ murine model not only has broad applications in COVID-19 vaccine studies, but also provides a unique rodent model for exploration of COVID-19-derived neuropathology. Based on the substantial permissibility of the brain to SARS-CoV-2 replication and development of a lethal disease, this pre-clinical model can be considered as a far more stringent than the golden hamster model.

The source of neurological manifestations associated with COVID-19 in patients with comorbid conditions can be: (i) direct impact of SARS-CoV-2 on CNS, (ii) infection of brain vascular endothelium and, (iii) uncontrolled anti-viral immune reaction inside CNS. ACE2 is expressed in human neurons, astrocytes and oligodendrocytes, located in middle temporal gyrus and posterior cingulate cortex, which may explain the brain permissibility to SARS-CoV-2 in patients (Song et al., 2020). Previous reports have demonstrated that respiratory viruses can invade the brain through neural dissemination or hematogenous route (Desforges et al., 2014). Besides that, the direct connection of olfactory system to the CNS via frontal cortex also represents a plausible route to invade the brain (Mori et al., 2005). Neural transmission of viruses to the CNS can occur as a result of direct neuron invasion through axonal transport in the olfactory mucosa. Subsequent to intraneuronal replication, the virus spreads to synapses and disseminate to anatomical CNS zones receiving olfactory tract projections (Koyuncu et al., 2013; Zubair et al., 2020; Berth, 2009; Koyuncu et al., 2013; Román et al., 2020). However, the detection of viral RNA in CNS regions without connection with olfactory mucosa suggests existence of another viral entry into the CNS, including migration of SARS-CoV-2-infected immune cells crossing the hemato-encephalic barrier or direct viral entry pathway via CNS vascular endothelium (Meinhardt et al., 2020). Although at steady state, viruses cannot penetrate to the brain through an intact blood-brain barrier (Berth, 2009), inflammation mediators which are massively produced during cytokine/chemokine storm, notably TNF-α and CCL2, can disrupt the integrity of blood-brain barrier or increase its permeability, allowing paracellular blood-to-brain transport of the virus or virus-infected leukocytes (Aghagoli et al., 2020; Hu et al., 2011).

The use of the highly stringent B6.K18-hACE2^IP-THV^ mice demonstrated the importance of i.n. booster immunization for inducing sterilizing protection of CNS by the LV-based vaccine candidate developed against SARS-CoV-2. Olfactory bulb may control viral CNS infection through the action of local innate and adaptive immunity (Durrant et al., 2016). In line with these observations, we detected increased frequencies of CD8^+^ T cells at this anatomically strategic area in i.m.-i.n. vaccinated and protected mice. In addition, substantial reduction in the inflammatory mediators was also found in the brain of the i.m.-i.n. vaccinated and protected mice, as well as decreased proportions of neutrophils and inflammatory monocytes respectively in the olfactory bulbs and brain. Regardless of the mechanism of the SARS-CoV-2 entry to the brain, we provide evidence of the full protection of the CNS against SARS-CoV-2 by i.n. booster immunization with NILV:: S_ΔF2P_.

We recently demonstrated the strong prophylactic capacity of LV::S_FL_ at inducing sterilizing protection in the lungs against SARS-CoV-2 infection (Ku et al., 2021). In the present study, moving toward a human clinical trial, we used LV encoding stabilized prefusion S_ΔF2P_ forms of S_Cov-2_. The choice of S_ΔF2P_ in this study was based on data indicating that stabilization of viral envelop glycoproteins at their prefusion forms improve the yield of their production as recombinant proteins in industrial manufacturing of subunit vaccines, and the efficacy of nucleic acid-based vaccines by raising availability of the antigen under its optimal immunogenic shape (Hsieh et al., 2020). The prefusion stabilization approach has been so far applied to S protein of several coronaviruses, including HKU1-CoV, SARS-CoV, and MERS-CoV. Stabilized S_MERS-CoV_ has been shown to elicit much higher NAb responses and protection in pre-clinical animal models (Hsieh et al., 2020). We detected no difference between the capacity of S_FL_ and S_ΔF2P_ at inducing anti-S_CoV-2_ IgG or IgA Ab responses in the sera or lung homogenates of LV-immunized animals. However, the possibility that the yield of LV::S_ΔF2P_ production at industrial level is higher is likely.

The sterilizing protection of the lungs conferred by a single i.n. administration and the full protection of CNS conferred by i.n. boost is an asset of primary importance. The non-cytopathic and non-inflammatory LV encoding either full length, or stabilized forms of S_CoV-2_, from either ancestral or emerging variants of SARS-CoV-2 provides a promising COVID-19 vaccine candidate of second generation. Protection of the brain, so far not directly addressed by other vaccine strategies, has to be taken into the account, considering the multiple and sometimes severe neuropathological manifestations associated with COVID-19.

## Acknowledgments

The authors are grateful to Pr Sylvie van der Werf (National Reference Centre for Respiratory Viruses hosted by Institut Pasteur, Paris, France) for providing the BetaCoV/France/IDF0372/2020 SARS-CoV-2 clinical isolate. The strain BetaCoV/France/IDF0372/2020 was supplied through the European Virus Archive goes Global (Evag) platform, a project that has received funding from the European Union’s Horizon 2020 research and innovation program under grant agreement No 653316. The authors thank Pr Geneviève Milon for fruitful advice and discussion, Dr Hugo Mouquet and Dr Cyril Planchais for providing recombinant homotrimeric S proteins, Dr Nicolas Escriou and Dr Marion Gransagne for providing a plasmid containing the S_ΔF2P_ sequence, Dr Jean Jaubert and Dr Xavier Montagutelli for the kind gift of B6.K18-ACE2^2Prlmn/JAX^ mice, Delphine Cussigh for her precious help with transgenic mice and Magali Tichit for excellent technical assistance in preparing histological sections.

This work was supported by the «URGENCE COVID-19» fundraising campaign of Institut Pasteur, TheraVectys and Agence Nationale de la Recherche (ANR) HuMoCID. Min Wen Ku is part of the Pasteur - Paris University (PPU) International PhD Program and received funding from the Institut Carnot Pasteur Microbes & Santé, and the European Union’s Horizon 2020 research and innovation program under the Marie Sklodowska-Curie grant agreement No 665807.

## Author Contribution

Study concept and design: MWK, MB, FA, FLV, LM, PC, acquisition of data: MWK, PA, MB, FA, AN, BV, FN, JL, PS, CB, KN, LM, construction and production of LV and technical support: PA, AN, FM, CB, PS, analysis and interpretation of data: MWK, PA, MB, FA, LM, PC, mouse transgenesis: SC, IL, FLV, histology: DH, FG, drafting of the manuscript: MWK, LM, PC.

## Methods

### Construction and production of LV::S_ΔF2P_

A codon-optimized S_ΔF2P_ sequence (1-1262) (Table S1) was amplified from pMK-RQ_S-2019-nCoV and inserted into pFlap by restriction/ligation between BamHI and XhoI sites, between the native human ieCMV promoter and a mutated Woodchuck Posttranscriptional Regulatory Element (WPRE) sequence. The *atg* starting codon of WPRE was mutated (mWPRE) to avoid transcription of the downstream truncated “X” protein of Woodchuck Hepatitis Virus for safety concerns (Figure S4). Plasmids were amplified and used to produce LV as previously described (Ku et al., 2021).

### Hamsters

Male *Mesocricetus auratus* golden hamsters (Janvier, Le Genest Saint Isle, France) were purchased mature and weighed between 100 to 120 gr at the beginning of the experiments. Hamsters were housed in individually-ventilated cages under specific pathogen-free conditions during the immunization period. For SARS-CoV-2 infection they were transferred into individually filtered cages placed inside isolators in the animal facility of Institut Pasteur. Prior to i.m. or i.n. injections, hamsters were anesthetized by isoflurane inhalation or i.p. injection of Ketamine (Imalgene, 80 mg/kg) and Xylazine (Rompun, 5 mg/kg).

### Mice

Female C57BL/6JRj mice (Janvier, Le Genest Saint Isle, France) were used between the age of 7 and 12 wks. Transgenic B6.K18-ACE2^2Prlmn/JAX^ mice (JAX stock #034860) were from Jackson Laboratories and were a kind gift of Dr Jean Jaubert and Dr Xavier Montagutelli (Institut Pasteur). Transgenic B6.K18-hACE2^IP-THV^ mice were generated and bred, as detailed below at the CIGM of Institut Pasteur. During the immunization period female or male transgenic mice were housed in individually-ventilated cages under specific pathogen-free conditions. Mice were transferred into individually filtered cages in isolator for SARS-CoV-2 inoculation at the Institut Pasteur animal facilities. Prior to i.n. injections, mice were anesthetized by i.p. injection of Ketamine (Imalgene, 80 mg/kg) and Xylazine (Rompun, 5 mg/kg).

### Mouse Transgenesis

The human K18 promoter (GenBank: AF179904.1 nucleotide 90 to 2579) was amplified by nested PCR from A549 cell lysate, as described previously (Chow et al., 1997; Koehler et al., 2000). The “i6×7” intron (GenBank: AF179904.1 nucleotide 2988 to 3740) was synthesized by Genscript. The K18^JAX^ (originally named K18i6×7PA) promoter includes the K18 promoter, the i6×7 intron at 5’ and an enhancer/polyadenylation sequence (PA) at 3’ of the *hACE2* gene. The^K18 IP-ThV^ promoter, instead of PA, contains the stronger wild-type WPRE element at 3 ’ of the *hACE2* gene. In contrast to K18^JAX^ construct which harbors the 3’ regulatory region containing a polyA sequence, the K18^IP-ThV^ construct takes benefice of the polyA sequence already present within the 3’ Long Terminal Repeats (LTR) of the lentiviral plasmid.

The i6×7 intronic part was modified to introduce a consensus 5’ splicing donor and a 3’ donor site sequence. The AAGGGG donor site was further modified for the AAGTGG consensus site. Based on a consensus sequence logo (Dogan et al., 2007), the poly-pyrimidine tract preceding splicing acceptor site (TACAATCCCTC in original sequence GenBank: AF179904.1 and TTTTTTTTTTT in K18^JAX^) was replaced by CTTTTTCCTTCC to limit incompatibility with the reverse transcription step during transduction. Moreover, original splicing acceptor site CAGAT was modified to correspond to the consensus sequence CAGGT. As a construction facility, a ClaI restriction site was introduced between the promoter and the intron. The construct was inserted into a pFLAP plasmid between the MluI and BamHI sites. h*ACE2* gene cDNA was introduced between the BamHI and XhoI sites by restriction/ligation. Integrative LV::K18-hACE2 was produced as described in (Ku et al., 2021) and concentrated by two cycles of ultracentrifugation at 22,000 rpm 1h 4°C.

ILV of high titre (4.16 × 10^9^ TU/ml) carrying K18-hACE2^IP-THV^ was used in transgenesis by subzonal micro-injection under the pellucida of fertilized eggs, and transplantation into the pseudo-pregnant B6CBAF1 females. LV allows particularly efficient transfer of the transgene into the nuclei of the fertilized eggs (Nakagawa and Hoogenraad, 2011). At N0 generation, ≈ 11% of the mice, i.e., 15 out of 139, had at least one copy of the transgene per genome. Eight N0 *hACE2^+^* males were crossed with female WT C57BL/6 mice. At N1 generation, ≈ 62% of the mice, i.e., 91 out of 147, had at least one copy of the transgene per genome.

### Genotyping and quantitation of *hACE2* gene copy number/genome in transgenic mice

Genomic DNA (gDNA) from transgenic mice was prepared from the tail biopsies by phenol-chloroform extraction. Sixty ng of gDNA were used as a template of qPCR with SYBR Green using specific primers listed in Table S2. Using the same template and in the similar reaction plate, mouse *pkdl* (Polycystic Kidney Disease 1) and *gapdh* were also quantified. All samples were run in quadruplicate in 10 μl reaction as follows: 10 min at 95°C, 40 cycles of 15 s at 95°C and 30 sec at 60°C. To calculate the transgene copy number, the 2^-ΔΔCt^ method was applied using the *pkd1* as a calibrator and *gapdh* as an endogenous control. The 2^-ΔΔCt^ provides the fold change in copy number of the *hACE2 gene* relative to *pkd1* gene.

### Ethical Approval of Animal Studies

Experimentation on mice and hamsters was realized in accordance with the European and French guidelines (Directive 86/609/CEE and Decree 87-848 of 19 October 1987) subsequent to approval by the Institut Pasteur Safety, Animal Care and Use Committee, protocol agreement delivered by local ethical committee (CETEA #DAP20007, CETEA #DAP200058) and Ministry of High Education and Research APAFIS#24627-2020031117362508 v1, APAFIS#28755-2020122110238379 v1.

### Humoral and T-cell immunity, Inflammation

As recently detailed elsewhere (Ku et al., 2021), T-splenocyte responses were quantitated by IFN-g ELISPOT and anti-S IgG or IgA Abs were detected by ELISA by use of recombinant stabilized S_CoV-2_. NAb quantitation was performed by use of S_CoV-2_ pseudo-typed LV, as previously described (Anna et al., 2020; Sterlin et al., 2020). The qRT-PCR quantification of inflammatory mediators in the lungs and brain of hamsters and mice was performed in total RNA extracted by TRIzol reagent, as recently detailed (Ku et al., 2021). CCL19 and CCL21 expression were measured using the following primer pairs: forward primers were 5’-CTG CCT CAG ATT ATC TGC CAT-3’ for CCL19 and 5’-AAG GCA GTG ATG GAG GGG-3’ for CCL21; reverse primers were 5’-AGG TAG CGG AAG GCT TTC AC-3’ for CCL19 and 5’-CGG GGT AAG AAC AGG ATT G-3’ for CCL21.

### SARS-CoV-2 inoculation

Hamsters or transgenic B6.K18-hACE2^IP-THV^ or B6.K18-ACE2^2Prlmn/JAX^ were anesthetized by i.p. injection of Ketamine and Xylazine mixture, transferred into a biosafety cabinet 3 and inoculated i.n. with 0.3 × 10^5^ TCID50 of the BetaCoV/France/IDF0372/2020 SARS-CoV-2 clinical isolate (Lescure et al., 2020). This clinical isolate was a gift of the National Reference Centre for Respiratory Viruses hosted by Institut Pasteur (Paris, France), headed by Pr. van der Werf. The human sample from which this strain was isolated has been provided by Dr. Lescure and Pr. Yazdanpanah from the Bichat Hospital, Paris, France. Mice or hamsters were inoculated i.n. with 20 μl or 50 μl of viral inoculum, respectively. Animals were housed in an isolator in BioSafety Level 3 animal facilities of Institut Pasteur. The organs recovered from the infected animals were manipulated according to the approved standard procedures of these facilities.

### Determination of viral loads in the organs

Organs from mice or hamsters were removed aseptically and immediately frozen at −80°C. RNA from circulating SARS-CoV-2 was prepared from lungs as recently described (Ku et al., 2021). Briefly, lung homogenates were prepared by thawing and homogenizing of the organs using lysing matrix M (MP Biomedical) in 500 μl of ice-cold PBS in an MP Biomedical Fastprep 24 Tissue Homogenizer. RNA was extracted from the supernatants of lung homogenates centrifuged during 10 min at 2000g. These RNA preparations were used to determine viral loads by E-specific qRT-PCR.

Alternatively, total RNA was prepared from lungs or other organs by addition of lysing matrix D (MP Biomedical) containing 1 mL of TRIzol reagent and homogenization at 30 s at 6.0 m/s twice using MP Biomedical Fastprep 24 Tissue Homogenizer. Total RNA was extracted using TRIzol reagent (ThermoFisher). These RNA preparations were used to determine viral loads by Esg-specific qRT-PCR, hACE2 expression level or inflammatory mediators.

SARS-CoV-2 E gene (Corman et al., 2020) or E sub-genomic mRNA (Esg mRNA) (Wolfel et al., 2020), was quantitated following reverse transcription and real-time quantitative TaqMan^®^ PCR, using SuperScriptTM III Platinum One-Step qRT-PCR System (Invitrogen) and specific primers and probe (Eurofins) (Table S3). The standard curve of Esg mRNA assay was performed using in vitro transcribed RNA derived from PCR fragment of “T7 SARS-CoV-2 Esg mRNA”. The in vitro transcribed RNA was synthesized using T7 RiboMAX Express Large Scale RNA production system (Promega) and purified by phenol/chloroform extraction and two successive precipitations with isopropanol and ethanol. Concentration of RNA was determined by optical density measurement, diluted to 10^9^ genome equivalents/μL in RNAse-free water containing 100μg/mL tRNA carrier, and stored at −80°C. Serial dilutions of this in vitro transcribed RNA were prepared in RNAse-free water containing 10μg/ml tRNA carrier to build a standard curve for each assay. PCR conditions were: (i) reverse transcription at 55°C for 10 min, (ii) enzyme inactivation at 95°C for 3 min, and (iii) 45 cycles of denaturation/amplification at 95°C for 15 s, 58°C for 30 s. PCR products were analyzed on an ABI 7500 Fast real-time PCR system (Applied Biosystems).

### Cytometric analysis of immune lung and brain cells

Isolation and staining of lung innate immune cells were largely detailed recently (Ku et al., 2021). Cervical lymph nodes, olfactory bulb and brain from each group of mice were pooled and treated with 400 U/ml type IV collagenase and DNase I (Roche) for a 30-minute incubation at 37°C. Cervical lymph nodes and olfactory bulbs were then homogenized with glass homogenizer while brains were homogenized by use of GentleMacs (Miltenyi Biotech). Cell suspensions were then filtered through 100 μm-pore filters, washed and centrifuged at 1200 rpm during 8 minutes. Cell suspensions from brain were enriched in immune cells on Percoll gradient after 25 min centrifugation at 1360 g at RT. The recovered cells from lungs were stained as recently described elsewhere (Ku et al., 2021). The recovered cells from brain were stained by appropriate mAb mixture as follows. (i) To detect innate immune cells: Near IR Live/Dead (Invitrogen), FcγII/III receptor blocking anti-CD16/CD32 (BD Biosciences), BV605-anti-CD45 (BD Biosciences), PE-anti-CD11b (eBioscience), PE-Cy7-antiCD11c (eBioscience), (ii) to detect NK, neutrophils, Ly-6C^+/-^ monocytes and macrophages: Near IR DL (Invitrogen), FcγII/III receptor blocking anti-CD16/CD32 (BD Biosciences), BV605-anti-CD45 (BD Biosciences), PE-anti-CD11b (eBioscience), PE-Cy7-antiCD11c (eBioscience), APC-anti-Ly6G (Miltenyi), BV711-anti-Siglec-F (BD), AF700-anti-NKp46 (BD Biosciences), FITC-anti-Ly6C (ab25025, Abcam) (iii) To detect adaptive immune cells: Near IR Live/Dead (Invitrogen), FcγII/III receptor blocking anti-CD16/CD32 (BD Biosciences), APC-anti-CD45 (BD), PerCP-Cy5.5-anti-CD3 (eBioscience), FITC-anti-CD4 (BD Pharmingen), BV711-anti-CD8 (BD Horizon), BV605-anti-CD69 (Biolegend), PE-anti-CCR7 (eBioscience) and VioBlue-Anti-B220 (Miltenyi). Cells were incubated with appropriate mixtures for 25 minutes at 4°C, washed in PBS containing 3% FCS and fixed with Paraformaldehyde 4% by an overnight incubation at 4°C. Samples were acquired in an Attune NxT cytometer (Invitrogen) and data analyzed by FlowJo software (Treestar, OR, USA).

### Lung Histopathology

Samples from the lungs or brain of hamsters or transgenic mice were fixed in formalin for 7 days and embedded in paraffin. Paraffin sections (5-μm thick) were stained with Hematoxylin and Eosin (H&E). Histopathological lesions were qualitatively described and when possible scored, using: (i) distribution qualifiers (i.e., focal, multifocal, locally extensive or diffuse), and (ii) a five-scale severity grade, i.e., 1: minimal, 2: mild, 3: moderate, 4: marked and 5: severe.

