## Supplemental Information for "Full Brain and Lung Prophylaxis against SARS-CoV-2 by Intranasal Lentiviral Vaccination in a New hACE2 Transgenic Mouse Model or Golden Hamsters"

**Supplemental Figures**

**Supplemental Tables**

#### Figure S1

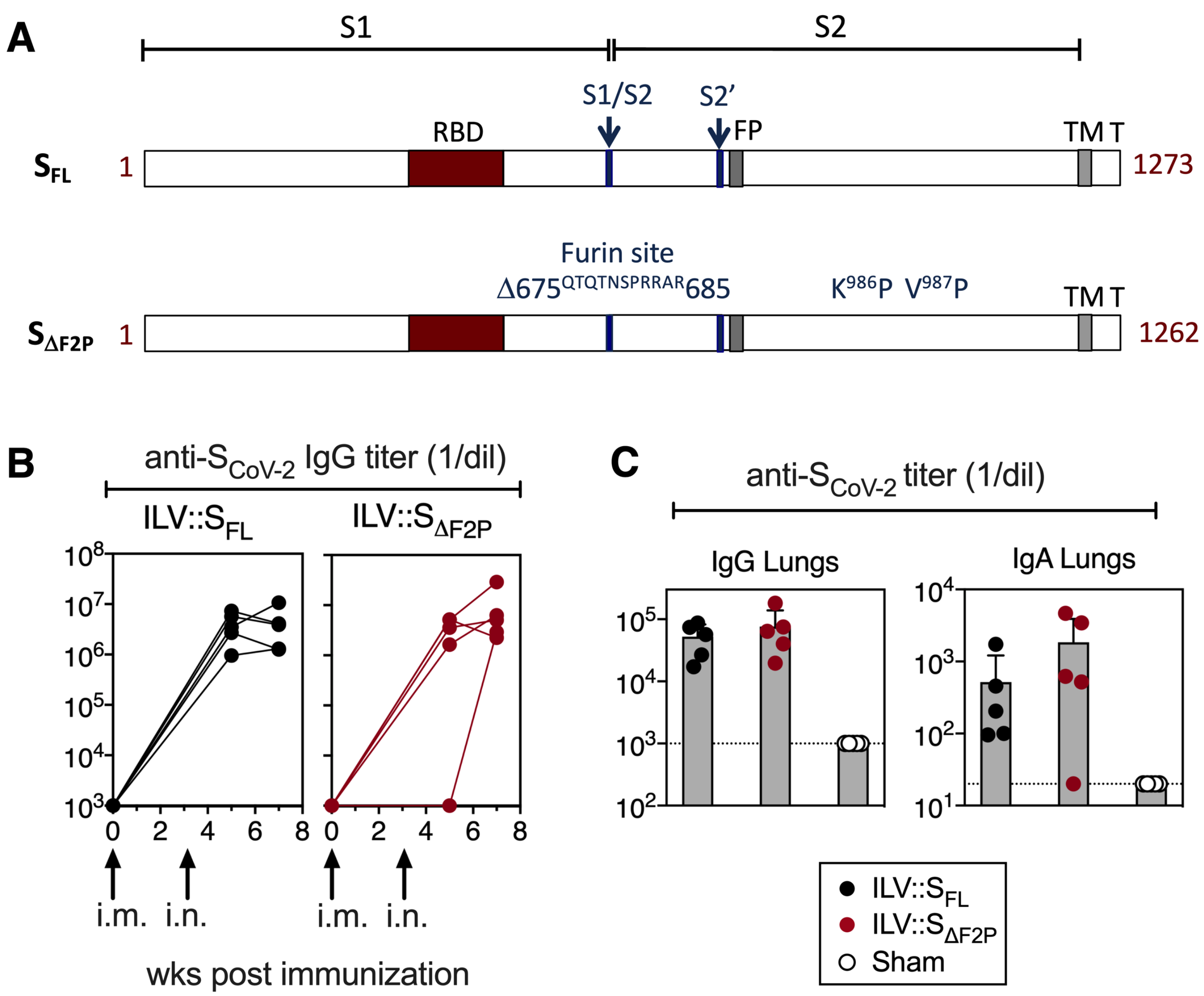


**Figure S1. Prefusion $S_{ΔF2P}$ protein and comparative adaptive immune responses induced following immunization of mice with LV::$S_{FL}$ or LV::$S_{ΔF2P}$. (A)** Schematic representation of $S_{FL}$ and $S_{ΔF2P}$ encoded by LV. RBD, S1/S2 and S2' cleavage sites, Fusion Peptide (FP), TransMembrane domain (TM) and short internal tail (T), $675^{QTQTNSPRRAR}685$ sequence encompassing RRAR furin cleavage site, and $K^{986}P$ and $V^{987}P$ consecutive substitutions are indicated. **(B-C)** C57BL/6 mice were primed i.m. at wk0 with $1 \times 10^7$ TU and boosted i.n. at wk5 with $3 \times 10^7$ TU of LV::$S_{FL}$, LV::$S_{ΔF2P}$ or a control LV (sham). **(B)** Sera were collected at 3, 5 and 7 wks post immunization and anti-$S_{CoV-2}$ (TriS) IgG responses were evaluated by ELISA. Results are expressed as mean endpoint dilution titers. **(C)** Lung homogenates were studied at 7 wks post immunization for anti-$S_{CoV-2}$ (TriS) IgG or IgA responses.

#### Figure S2

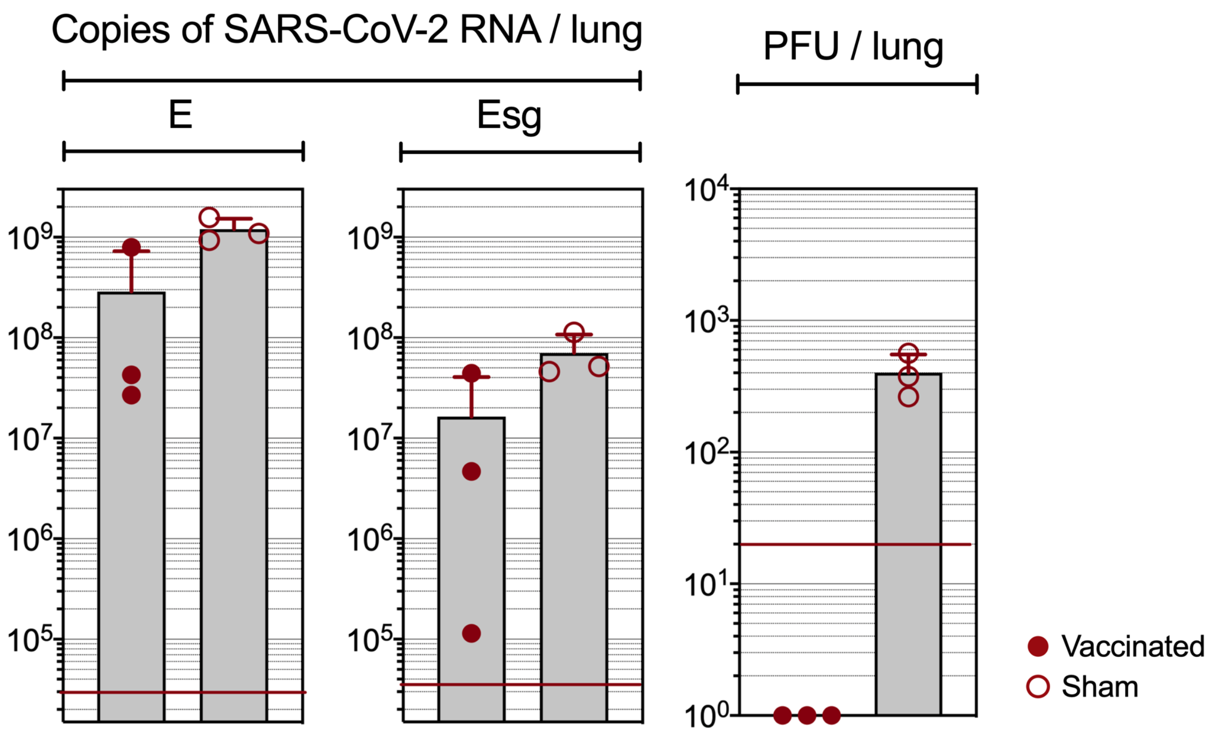


**Figure S2. Quantitation of lung SARS-CoV-2 loads PFU assay can be biased in vaccinated animals.** Lung viral loads in vaccinated or unvaccinated controls, as determined by conventional E-specific or sub-genomic Esg-specific qRT-PCR or by PFU counting. Note that vaccinated animals can have high loads which are not detectable by PFU, which can be explained by the presence of NAbs in the lungs which bias quantitation of cultivable viral particles.

#### Figure S3

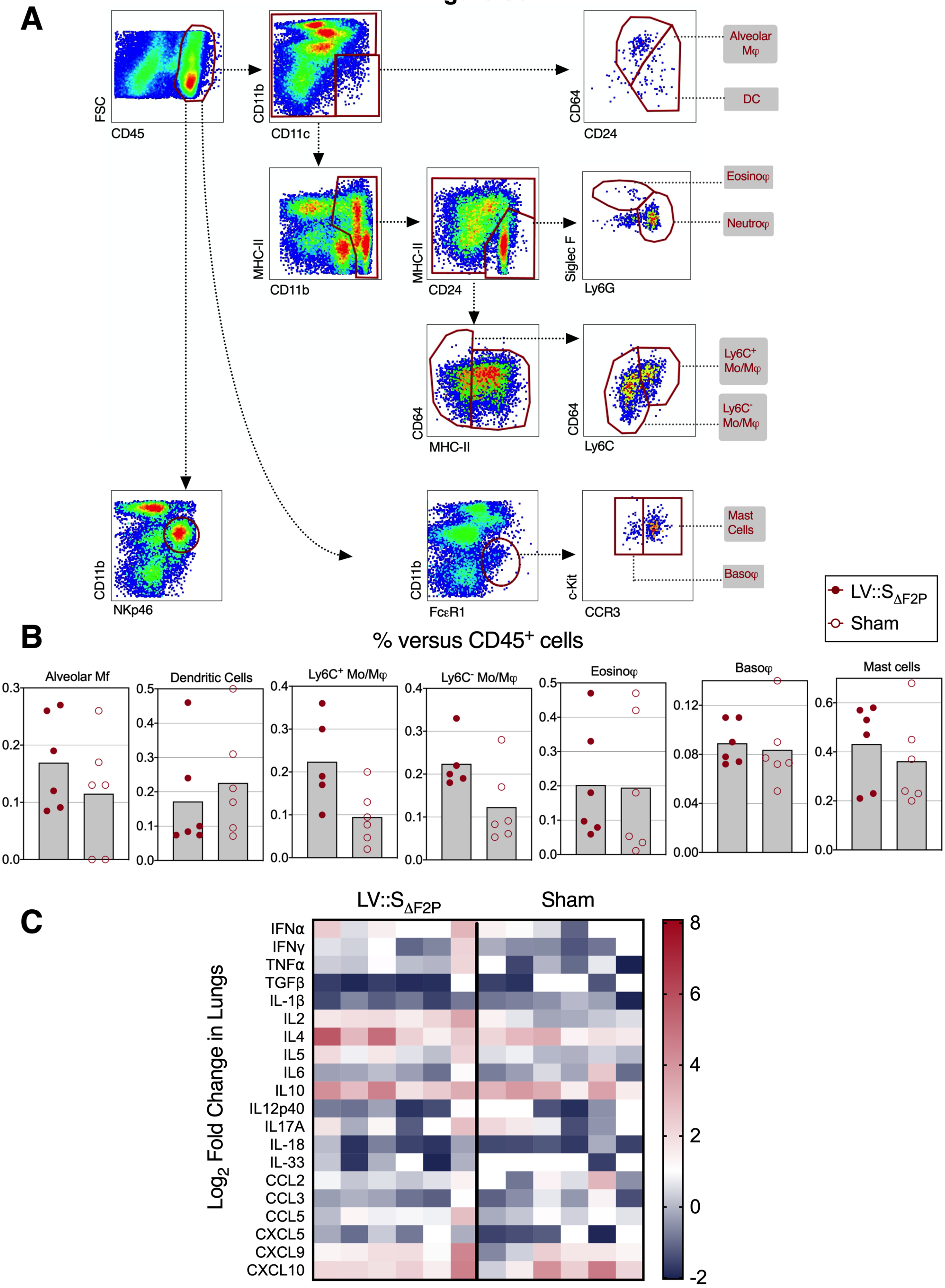


**Figure S3. Cytometric analysis of innate cell population in the lungs of LV::S$_{\Delta F2P}$- or sham-vaccinated and SARS-CoV-2-challenged K18-hACE2$^{IP-THV}$ transgenic mice. (A)** Cytometric gating strategy to quantify various lung innate immune cells at 3 dpi. Cells were first gated on hematopoietic CD45$^{+}$ cells and then by sequential gates, through three distinct paths. **(B)** Percentages of each innate immune subset were calculated versus total lung CD45$^{+}$ cells. **(C)** Heatmap recapitulating $\log_2$ fold change in cytokine and chemokine mRNA expression in the lungs of B6.K18-hACE2$^{IP-THV}$ and B6.K18-ACE2$^{2Prlmn/JAX}$ transgenic mice at 3 dpi. Data were normalized versus untreated controls.

### Figure S4

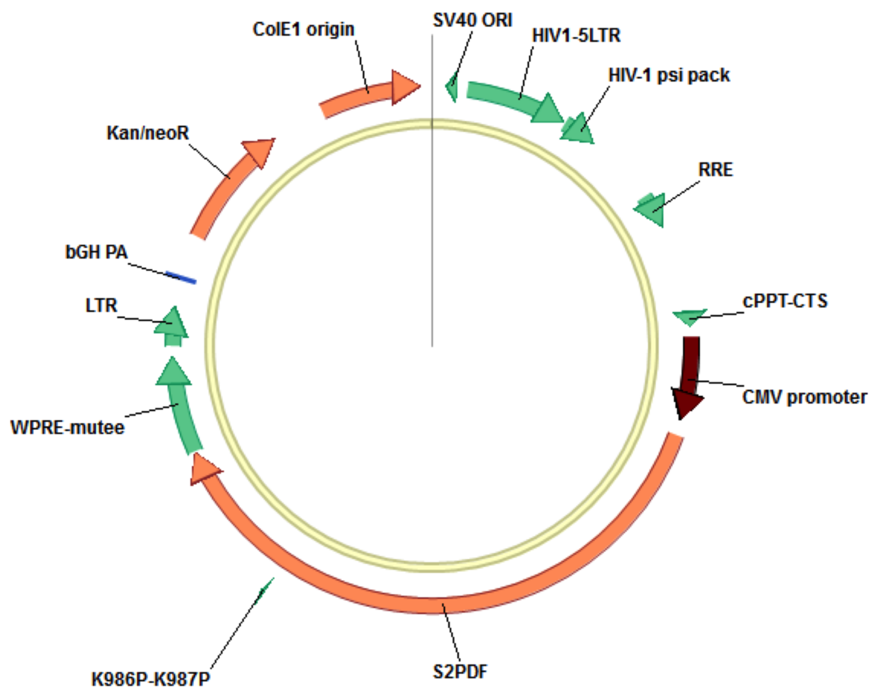


**pFlap-CMV-S$_{\Delta F2P}$-WPREm**

10046 bp

**Figure S4. Maps of lentiviral plasmid encoding for S$_{\Delta F2P}$**

**Table S1. Sequences of prefusion $S_{\Delta F2P}$, as encoded by ILV or NILV vaccinal vectors**

| **Pre-fusion** | **a.a. sequence** |
|---|---|
| $S_{\Delta F2P}$ | MFVFLVLLPLVSSQCVNLTTRTQLPPAYTNSFTRGVYYPDKVFRSSVLHSTQDLFLPFFSNVTWFH<br>AIHVSGTNGTKRFDNPVLPFNDGVYFASTEKSNIIRGWIFGTTLDSKTQSLLIVNNATNVVIKVCEF<br>QFCNDPFLGVYYHKNNKSWMESEFRVYSSANNCTFEYVSQPFLMDLEGKQGNFKNLREFVFKNI<br>DGYFKIYSKHTPINLVRDLPQGFSALEPLVDLPIGINITRFQTLLALHRSYLTPGDSSSGWTAGAAA<br>YYVGYLQPRTFLLKYNENGTITDAVDCALDPLSETKCTLKSFTVEKGIYQTSNFRVQPTESIVRFP<br>NITNLCPFGEVFNATRFASVYAWNRKRISNCVADYSVLYNSASFSTFKCYGVSPTKLNDLCFTNV<br>YADSFVIRGDEVRQIAPGQTGKIADYNYKLPDDFTGCVIAWNSNNLDSKVGGNYNYLYRLFRKS<br>NLKPFERDISTEIYQAGSTPCNGVEGFNCYFPLQSYGFQPTNGVGYQPYRVVVLSFELLHAPATVC<br>GPKKSTNLVKNKCVNFNFNGLTGTGVLTESNKKFLPFQQFGRDIADTTDAVRDPQTLEILDITPCS<br>FGGVSVITPGTNTSNQVAVLYQDVNCTEVPVAIHADQLTPTWRVYSTGSNVFQTRAGCLIGAEHV<br>NNSYECDIPIGAGICASY**QTQTNSPRRAR**SVASQSIIAYTMSLGAENSVAYSNNSIAIPTNFTISVTT<br>EILPVSMTKTSVDCTMYICGDSTECSNLLLQYGSFCTQLNRALTGIAVEQDKNTQEVFAQVKQIY<br>KTPPIKDFGGFNFSQILPDPSKPSKRSFIEDLLFNKVTLADAGFIKQYGDCLGDIAARDLICAQKFN<br>GLTVLPPLLTDEMIAQYTSALLAGTITSGWTFGAGAALQIPFAMQMAYRFNGIGVTQNVLYENQK<br>LIANQFNSAIGKIQDSLSSTASALGKLQDVVNQNAQALNTLVKQLSSNFGAISSVLNDILSRLD**PP**E<br>AEVQIDRLITGRLQSLQTYVTQQLIRAAEIRASANLAATKMSECVLGQSKRVDFCGKGYHLMSFP<br>QSAPHGVVFLHVTYVPAQEKNFTTAPAICHDGKAHFPREGVFVSNGTHWFVTQRNFYEPQIITTD<br>NTFVSGNCDVVIGIVNNTVYDPLQPELDSFKEELDKYFKNHTSPDVDLGDISGINASVVNIQKEID<br>RLNEVAKNLNESLIDLQELGKYEQYIKWPWYIWLGFIAGLIAIVMVTIMLCCMTSCCSCLKGCCS<br>CGSCCKFDEDDSEPVLKGVKLHYT |

The deleted sequence encompassing the furin cleavage site and double proline substitution in S2 are indicated in red.

**Table S2. Sequences of primers used to genotype B6.K18-hACE2$^{IP-THV}$ transgenic mice.**

| **Primers** | |
|---|---|
| hACE2 Fw | TCCTAACCAGCCCCCTGTT |
| hACE2 Rv | TGACAATGCCAACCA CTATCACT |
| PKD1 Fw | GGCTGCTGAGCGTCTGGTA |
| PKD1 Rv | CCAGGTCCTGCGTGTCTGA |
| GAPDH-ACE2 Fw | GCCCAGAACATCATCCCTGC |
| GAPDH-ACE2 Rv | CCGTTCAGCTCTGGGATGACC |

**Table S3. Sequences of primers used to quantitate SARS-CoV-2 loads by qRT-PCR**

| Primer/Probe | DNA Sequence |
|---|---|
| “E-Sarbeco” Fw | 5’-ACAGGTACGTTAATAGTTAATAGCGT-3’ |
| “E-Sarbeco” Rv | 5’-ATATTGCAGCAGTACGCACACA-3’ |
| “E-Sarbeco” | 5’-FAM-ACACTAGCCATCCTTACTGCGCTTCG-BHQ-1-3’ |
| “Esg mRNA” Fw | 5’-CGATCTCTTGTAGATCTGTTCTC-3’ |
